# Human V-ATPase a-subunit isoforms bind specifically to distinct phosphoinositide phospholipids

**DOI:** 10.1101/2023.04.24.538068

**Authors:** Connie Mitra, Patricia M. Kane

## Abstract

V-ATPases are highly conserved multi-subunit enzymes that maintain the distinct pH of eukaryotic organelles. The integral membrane a-subunit is encoded by tissue and organelle specific isoforms, and its cytosolic N-terminal domain (aNT) modulates organelle specific regulation and targeting of V-ATPases. Organelle membranes have specific phosphatidylinositol phosphate (PIP) lipid enrichment linked to maintenance of organelle pH. In yeast, the aNT domains of the two a-subunit isoforms bind PIP lipids enriched in the organelle membranes where they reside; these interactions affect activity and regulatory properties of the V-ATPases containing each isoform. Humans have four a-subunit isoforms. We hypothesize that the aNT domains of the human isoforms will also bind to specific PIP lipids. The a1 and a2 isoforms of human V-ATPase a-subunits are localized to endolysosomes and Golgi, respectively. Bacterially expressed Hua1NT and Hua2NT bind specifically to endolysosomal PIP lipids PI(3)P and PI(3,5)P2 and Golgi enriched PI(4)P, respectively. Despite the lack of canonical PIP binding sites, potential binding sites in the HuaNT domains were identified by sequence comparisons and existing subunit structures and models. Mutations at a similar location in the distal loops of both HuaNT isoforms compromise binding to their cognate PIP lipids, suggesting that these loops encode PIP specificity of the a-subunit isoforms. These data also suggest a mechanism through which PIP lipid binding could stabilize and activate V-ATPases in distinct organelles.

## INTRODUCTION

The membrane bound organelles in eukaryotic cells each have a distinct, tightly regulated luminal pH important for organelle function and cell growth. This specific pH supports organelle identity and is established principally by vacuolar H+-ATPases (V-ATPases) (Maxson and Grinstein 2014, Banerjee and Kane 2017). V-ATPases are highly conserved multi-subunit enzymes, functioning as ATP-driven proton pumps (Collins and Forgac 2020). They are comprised of 14 subunits organized in two operational subcomplexes, V_1_ and V_o_ (Oot, Couoh-Cardel et al. 2017). The catalytic V_1_ subcomplex is a peripheral membrane complex on the cytosolic face of the membrane and the V_o_ subcomplex is embedded in the membrane (Kane 2006). The V_1_ subcomplex executes ATP hydrolysis and the V_o_ subcomplex contains the proton pore (Graham, Powell et al. 2000, Oot, Couoh-Cardel et al. 2017, Seidel 2022). V-ATPases are ubiquitously expressed and localized to multiple organelles, including the Golgi, endosomes, secretory vesicles, lysosomes and the lysosome-like yeast and plant vacuoles (Maxson and Grinstein 2014, Seidel 2022). Their principal function is organelle acidification and through this function they impact processes such as receptor mediated endocytosis, protein processing, trafficking and degradation, neurotransmitter loading, and modulation of signaling pathways such as Notch and mTOR (Maxson and Grinstein 2014, Pamarthy, Kulshrestha et al. 2018, Collins and Forgac 2020). In mammals, V-ATPases are also present in the plasma membrane of specific cells including kidney intercalated cells, epididymal clear cells, osteoclasts, macrophages and neutrophils. In these settings, they acidify the extracellular space and function in pH homeostasis, bone resorption, urine acidification and sperm maturation. (Breton and Brown 2013, Stransky, Cotter et al. 2016, Chu, Zirngibl et al. 2021)

The a-subunit of the V_o_ domain of V-ATPases is a 100 kDa subunit, which consists of amino terminal (aNT) and carboxy terminal (aCT) domains (Figure 1A). The aNT is on the cytosolic face of the membrane and hydrophilic and the aCT is integral to the membrane and hydrophobic (Oot, Couoh-Cardel et al. 2017, Roh, Stam et al. 2018). The aCT domain is involved in proton translocation (Kawasaki-Nishi, Bowers et al. 2001, Arata, Nishi et al. 2002, Qi and Forgac 2008). Cryo-EM studies establish that the aNT domain resembles a folded hairpin, comprised of globular proximal and distal ends connected by a coiled-coil linker, giving it a dumbbell shape (Roh, Stam et al. 2018). The aNT domain functions as a regulatory core for several reasons. It is positioned at the V_1_-V_o_ interface and is capable of interacting with the three peripheral stator stalks in the assembled enzyme. In intact, active V-ATPase, the aNT proximal and distal domains are pulled away from the central stalks because of their interactions with stator stalks (Oot and Wilkens 2012, Oot, Couoh-Cardel et al. 2017), while in autoinhibited V_o_ that is not bound to V_1_, the aNT domain collapses towards the central stalk (Roh, Stam et al. 2018). These conformational changes affect both the proximal and distal ends of aNT and may be involved in V-ATPase regulation by reversible disassembly (Stam and Wilkens 2017). In addition, aNT domains interact with cellular factors essential for V-ATPase regulation. These factors include RAVE, PIP lipids, and regulatory proteins like glycolytic enzymes (Banerjee and Kane 2020, Collins and Forgac 2020, Jaskolka, Winkley et al. 2021). Significantly, the information for compartment specific targeting, localization and signaling of V-ATPase to specific cellular membranes also is encoded in the aNT domain (Kawasaki-Nishi, Bowers et al. 2001, Banerjee and Kane 2017).

**Figure 1:**
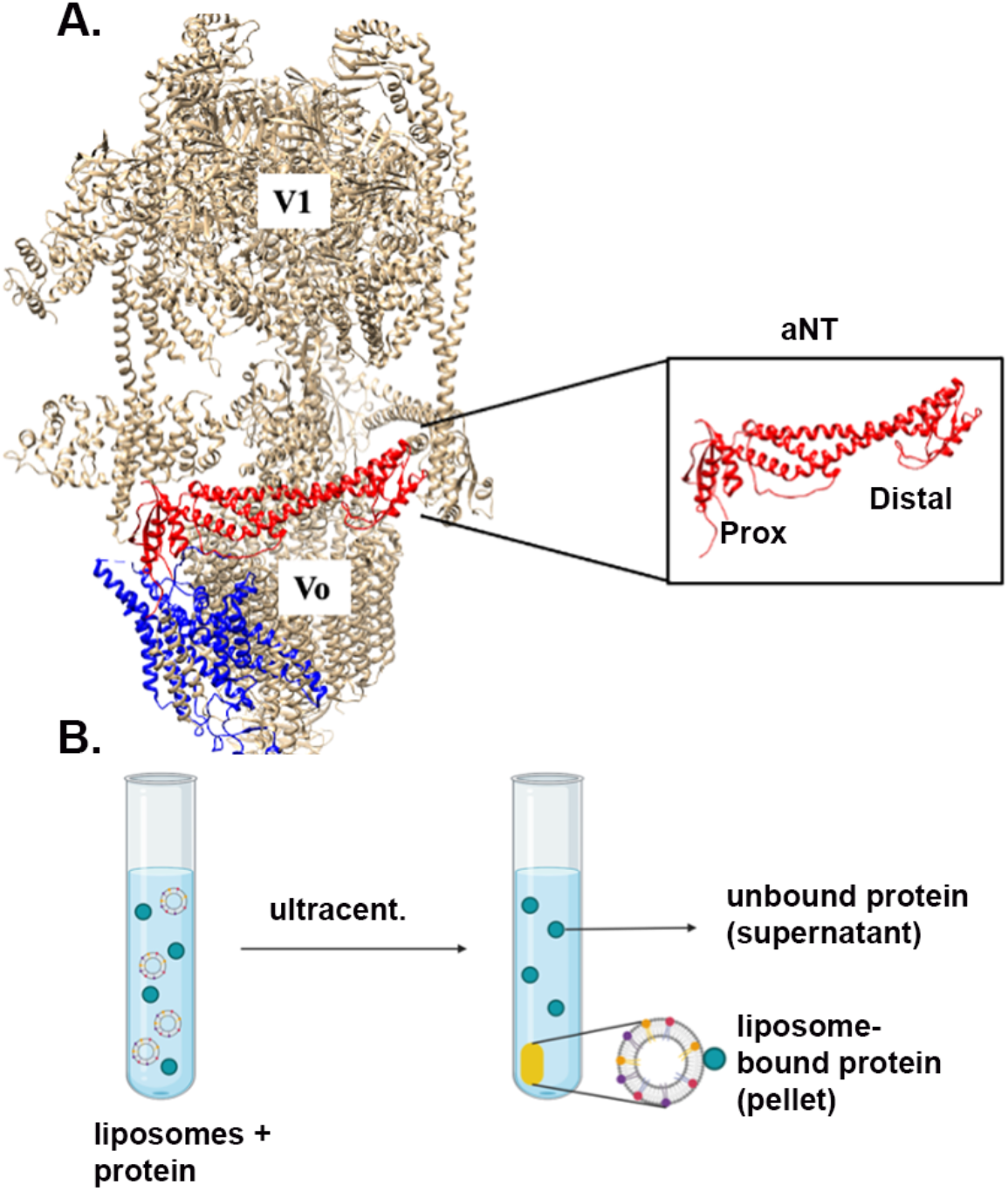
Human V-ATPase structure and liposome pelleting Assay. **A**. Cryo-EM structure of V-ATPase from human embryonic kidney cell line HEK293F (PDB 6WM2). The cytosolic and transmembrane sectors are denoted as V_1_ and V_o_. The V_o_ aNT domain is shown in red and aCT domain in blue. The structure of the a1NT is shown on the left, with the proximal and distal ends indicated. **B**. Overview of the liposome pelleting assay. Liposomes of defined lipid content are combined with varied protein concentrations, incubated, and subjected to ultracentrifugation as describe in Experimental Procedures. Liposomes bound to protein are in pellet and unbound protein are in supernatant. The image is created with BioRender.com.

Importantly, the V-ATPase a-subunits are expressed as multiple isoforms that are critical for targeting of V-ATPases to specific compartments and administer their manifold organelle-specific functions to V-ATPase (Nishi and Forgac 2000, Toyomura, Oka et al. 2000, Sun-Wada and Wada 2022). Yeast has two a-subunit isoforms, Vph1 and Stv1, which are localized at steady state to vacuoles and Golgi (Kane 2006, Toei, Saum et al. 2010). Higher organisms, including humans, have four isoforms designated a1-a4 (Forgac 2007). The a1, a2 and a3 isoforms are expressed ubiquitously but can also be enriched in certain organelles or cell types. The a1 isoform is present in lysosomes, but is also enriched in synaptic vesicles (Vasanthakumar and Rubinstein 2020). The a2 isoform is predominantly localized to the Golgi apparatus but has also been found in endosomes (Hurtado-Lorenzo, Skinner et al. 2006). The a3 isoform is present in lysosomes, particularly secretory lysosomes, phagosomes, and the plasma membrane of osteoclasts (Ochotny, Van Vliet et al. 2006, Sun-Wada, Tabata et al. 2009). The a4 isoform is specific to renal intercalated cells and epididymal clear cells (Breton and Brown 2013). Genetic defects in V-ATPase activity due to loss of specific subunit isoforms are implicated in several diseases, including distal renal tubule acidosis (a4 mutations) (Breton and Brown 2013), cutis laxa type II (a2 mutations) (Kornak, Reynders et al. 2008), and autosomal recessive osteopetrosis (a3 mutations) (Frattini, Orchard et al. 2000). Also, aberrant V-ATPase activity is implicated in cancer and viral infection (Kohio and Adamson 2013, Stransky, Cotter et al. 2016). This makes V-ATPase an attractive target for drug development.

Another important feature of eukaryotic organelle membranes is the enrichment of phosphatidylinositol phosphate (PIP) lipids. PIPs are transiently generated from phosphatidylinositol (PI) lipids in outer leaflets of cellular membrane by lipid kinases and phosphatases (Strahl and Thorner 2007). Their levels are scarce, but they can modulate vesicular trafficking, ion channels, pumps and transporters and maintain organelle physiology (Balla 2013, Hille, Dickson et al. 2015, De Craene, Bertazzi et al. 2017). In eukaryotic cells, PIP lipids are enriched in specific intracellular membranes where they help to specify organelle identity (Shewan, Eastburn et al. 2011, Ho, Alghamdi et al. 2012, Hammond and Balla 2015, De Craene, Bertazzi et al. 2017). PI(3)P is enriched in early endosomes, but are also present in late endosomes and lysosomes (Clague, Urbé et al. 2009, Shewan, Eastburn et al. 2011, Hammond and Balla 2015, Wallroth and Haucke 2018). PI(4)P is the chief PIP lipid in the Golgi apparatus but is found in plasma membranes as well (Shewan, Eastburn et al. 2011, Vasanthakumar and Rubinstein 2020). PI(3,5)P_2_ is predominantly localized in yeast vacuoles and mammalian late endosomes, lysosomes and multivesicular bodies (Strahl and Thorner 2007, Dove, Dong et al. 2009, Shewan, Eastburn et al. 2011, Ho, Alghamdi et al. 2012, McCartney, Zhang et al. 2014). PI(4,5)P_2_ is enriched in the plasma membrane (Hammond, Fischer et al. 2012).

V-ATPase activity is regulated by PIP lipids in yeast. V-ATPases containing the Vph1 a-subunit isoform are activated and stabilized by salt stress and high extracellular pH in a PI(3,5)P_2_-dependent manner (Li, Diakov et al. 2014). Addition of short chain PI(3,5)P_2_ to isolated vacuoles increases V-ATPase and proton pumping activity (Banerjee, Clapp et al. 2019). Mutations in the distal domain of Vph1NT compromise this *in vitro* activation, and when present in intact Vph1, also impact the cellular response to osmotic stress (Banerjee, Clapp et al. 2019). Elevating PI(3,5)P_2_ levels by constitutive activation of Fab1 kinase or salt shock reversibly recruits Vph1NT to membranes *in vivo* in the absence of other subunits (Li, Diakov et al. 2014). Taken together, these data suggest that Vph1NT can interact with PI(3,5)P_2_. PI(4)P is also essential for Stv1 localization and activity. *In vitro*, Stv1NT binds PI(4)P and this association is impaired by a mutation in the proximal domain of Stv1NT (Banerjee and Kane 2017). When this mutation is introduced into full length Stv1, or Golgi PI(4)P levels are lowered, Stv1-containing V-ATPases are mislocalized to the vacuole (Banerjee and Kane 2017). Addition of PI(4)P was shown to enhance activity of Stv1-containing V-ATPases (Vasanthakumar, Bueler et al. 2019). These data highlight the important of PIP interactions with V-ATPase a-subunit isoforms.

In this paper, we aim to determine whether PIP binding is conserved in mammalian a-subunit isoforms, and to elucidate any PIP specificity of the human a1 and a2 isoforms. We have tested the specificity of Hua1NT and Hua2NT for intracellular PIP lipids in a liposome pelleting assay and found that Hua1NT binds preferentially to liposomes containing PI(3)P and PI(3,5)P_2_, lipids typically enriched in endosomes and lysosomes. In contrast, Hua2NT shows a preference for Golgi-enriched lipid PI(4)P. We have identified potential PIP specific binding sites for both Hua1NT and Hua2NT in their distal domains. The data demonstrate the association between specific V-ATPase subunit isoforms and the PIP lipids enriched in the organelles where they reside. Defining PIP binding codes on V-ATPase will improve our understanding of organelle specific pH control and could provide new avenues for controlling V-ATPase subpopulations.

## RESULTS

### Hua1NT binds the endolysosomal PIP lipids PI(3)P and PI(3,5)P2 *in vitro*

The human a1 and a2 isoforms reside in endolysosomal compartments and Golgi, respectively, in most mammalian cells. Based on the results in yeast indicating that vacuolar Vph1NT binds to PI(3,5)P_2_ and Stv1NT binds to PI(4)P, we hypothesized that the human a1 and a2 subunits might also bind to the predominant intracellular PIP lipids in their organelles of residence, including PI(3)P, PI(3,5)P_2_ and PI(4)P. In addition, based on current structures and results with the yeast a-subunits, we hypothesized that the cytosolic N-terminal domains would be most likely to contain PIP lipid binding sites.

To investigate whether the N-terminal domain of human Vo a-subunit a1 (Hua1NT) and a2 (Hua2NT) isoforms exhibit specific binding to PIP lipids. We expressed the NT domain only of each isoform (Figure 1A) and tested their binding to liposomes in a quantitative liposome pelleting assay (Figure 1B) (Chandra, Chin et al. 2019). The Hua1NT domain (aa 1-356) was fused to maltose-binding protein (MBP) at its N-terminus and a FLAG tag at its C-terminus to generate MBP-Hua1NT-FLAG. The tagged protein was expressed in *Escherichia coli* and purified by sequential affinity chromatography on amylose and anti-FLAG columns, followed by size exclusion chromatography. Purity was analyzed by SDS-PAGE (Figure 2A, left). The Hua1NT construct eluted at the molecular mass of a monomer (M) from the gel filtration column (Figure 2A, right). This fraction was used to characterize the *in vitro* binding specificity of Hua1NT and different PIP lipids. In this assay, different concentrations of the expressed protein were mixed with liposomes containing 5 mol% of the indicated PIP lipid, then incubated at room temperature to allow interaction, followed by ultracentrifugation (described in Experimental Procedures). The same concentration of protein was centrifuged without any liposomes in order to determine whether there is any precipitation; this is the “Protein only” sample. As an additional control, we incubate the proteins with liposomes without PIP lipids, but with an equal amount of phosphatidyl serine (PS) replacing the PIPs. This sample serves as a control for liposome binding that is not PIP-specific. For all samples, supernatant and pellet fractions were obtained as described in Experimental Procedures and analyzed by SDS-PAGE and immunoblotting.

**Figure 2:**
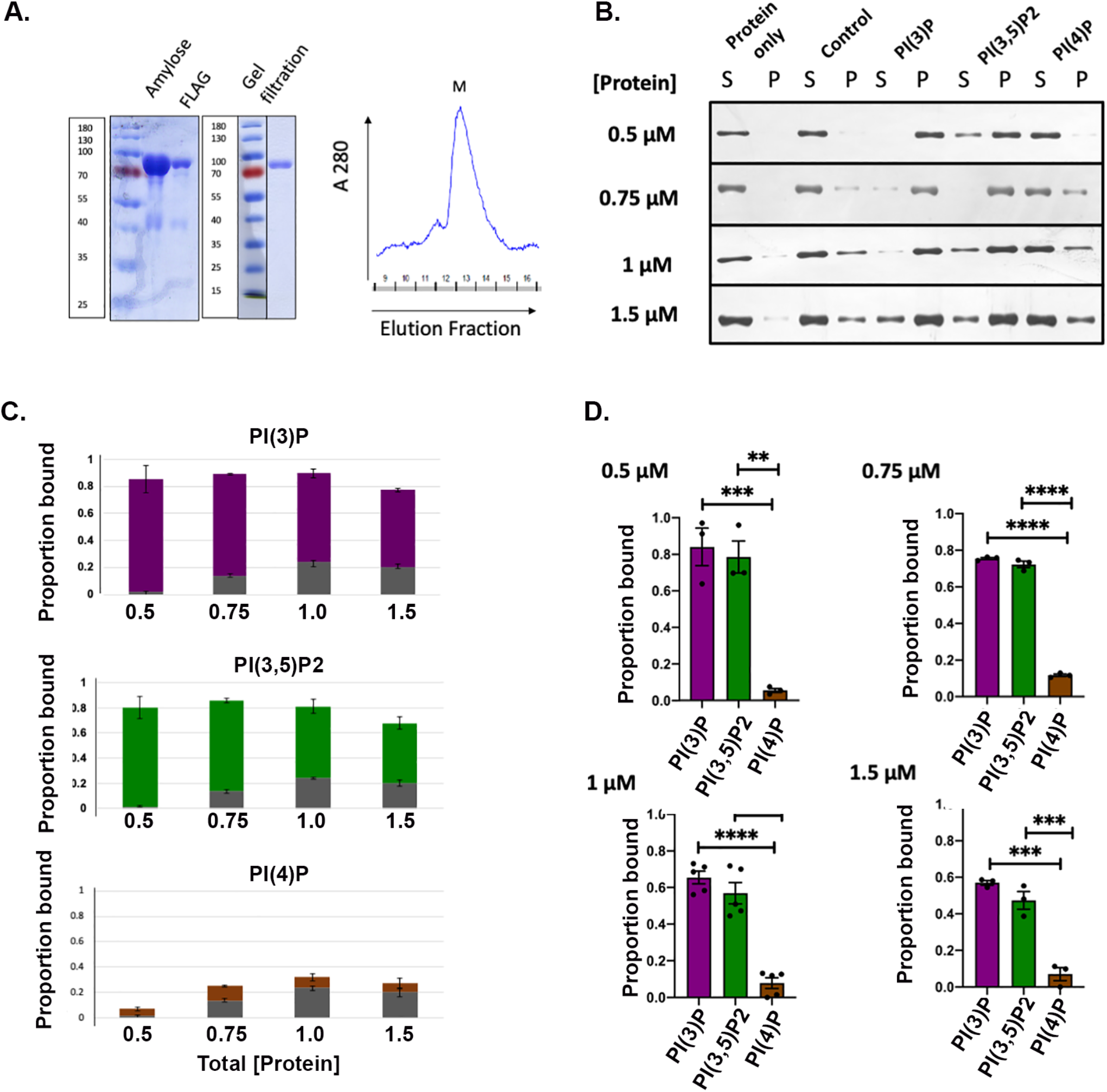
Hua1NT binds the endolysosomal PIP lipids PI(3)P and PI(3,5)P_2_ *in vitro*. **A**. Left, Coomassie-stained SDS-PAGE showing purified MBP-Hua1NT-FLAG after amylose, FLAG column and gel filtration purification, with the molecular mass markers indicated. Right, Elution profile of MBP-Hua1NT-FLAG from gel filtration column. The peak corresponding to the monomeric molecular mass (M) was collected and used for the liposome pelleting assay. **B**. Anti-FLAG immunoblot of MBP-Hua1NT-FLAG in supernatant (S) and Pellet (P) fractions collected after ultracentrifugation. Protein only samples contained no liposomes. Control samples used liposomes containing no PIP lipid. Experimental samples used liposomes containing 5% of the indicated PIP lipids. The experiments were performed using different protein concentrations as indicated to the left of the blots. **C**. Stacked plots showing average proportion (± S.E.M.) of protein bound to control and experimental samples. Control binding is indicated by gray area on the bottom of each stack, PIP-specific binding is indicated by the colored portion at the top. **D**. PIP-specific binding of MBP-Hua1NT-FLAG at different protein concentrations. The proportion of protein bound to control in **C**. was subtracted from the total to give the PIP-specific binding at each protein concentration. At least three assays were performed for each protein concentration and PIP lipid, and each dot represents a distinct assay (biological replicate). Bars represent mean ± S.E.M. Statistical significance was determined by ordinary one-way ANOVA with Tukey’s multiple comparison test. **** P<0.0001, *** P<0.0005, ** P<0.005, * P<0.05, ns P>0.05.

Hua1NT interacted with both PI(3)P and PI(3,5)P_2_ as indicated by the presence of bands in pellet (P) fraction at all concentrations (Figure 2B). In contrast, Hua1NT pelleted much less with PI(4)P liposomes, suggesting weaker binding (Figure 2B). The relative amount of protein in the supernatant and pellet fractions were determined by quantification of band intensity by ImageJ for each experiment and dividing the signal from the pellet over the total (supernatant + pellet) signal. The level of precipitation in the protein only sample was negligible. There was some variability in individual samples, but each experiment was repeated at least three times with freshly prepared protein and liposomes. The average proportion of total protein that bound to the PIP-containing and control liposomes over several experiments is plotted in Figure 2C. The control, PIP-independent binding (bottom of stacked bars for each protein concentration) is relatively low. In Figure 2D, PIP-independent binding is subtracted in order to compare the level of PIP-specific binding at each protein concentration. These data indicate that Hua1NT binds specifically PI(3)P and PI(3,5)P2 liposomes, but shows little binding to PI(4)P-containing liposomes. In addition, binding is nearly complete at the lowest concentration of protein measured (0.5 µM), indicating sub-micromolar affinity of the protein for these lipids. These data suggest that the Hua1 isoform, which resides in the endosomes and lysosomes of many cells, shows preferential binding to PI(3)P and PI(3,5)P_2_, the lipids enriched in these organelles.

### Hua2NT binds the Golgi PIP lipid PI(4)P *in vitro*

We next tested the specificity of Hua2NT for the same three PIP lipids. MBP-Hua2NT-FLAG (aa 1-364) was expressed in *Escherichia coli* and purified similarly to Hua1NT. A Coomassie stained SDS-PAGE of purified MBP-Hua2NT-FLAG after affinity purification and gel filtration and its elution profile are depicted in Figure 3A. The liposome pelleting assay was performed to test the interaction of Hua2NT with liposomes of distinct PIP content. We found that at different concentrations, Hua2NT interacted with PI(4)P most strongly, as indicated by the bands in pellet (P) fractions (Figure 3B). There is again negligible pelleting in the protein only sample. The pelleting of Hua2NT with PI(3)P and PI(3,5)P_2_ was less than that with PI(4)P (Figure 3B). The proportion of protein at different concentrations bound to the PIP lipids and control was determined as described above and is plotted in Figure 3C. The binding to control liposomes is relatively low (Figure 3C) and was subtracted in Figure 3D. The comparison indicates Hua2NT binds specifically and with relatively high affinity to PI(4)P (Figure 3D).These data suggest that the Hua2 isoform, which functions in Golgi, shows a preference for the Golgi-enriched PI(4)P.

**Figure 3:**
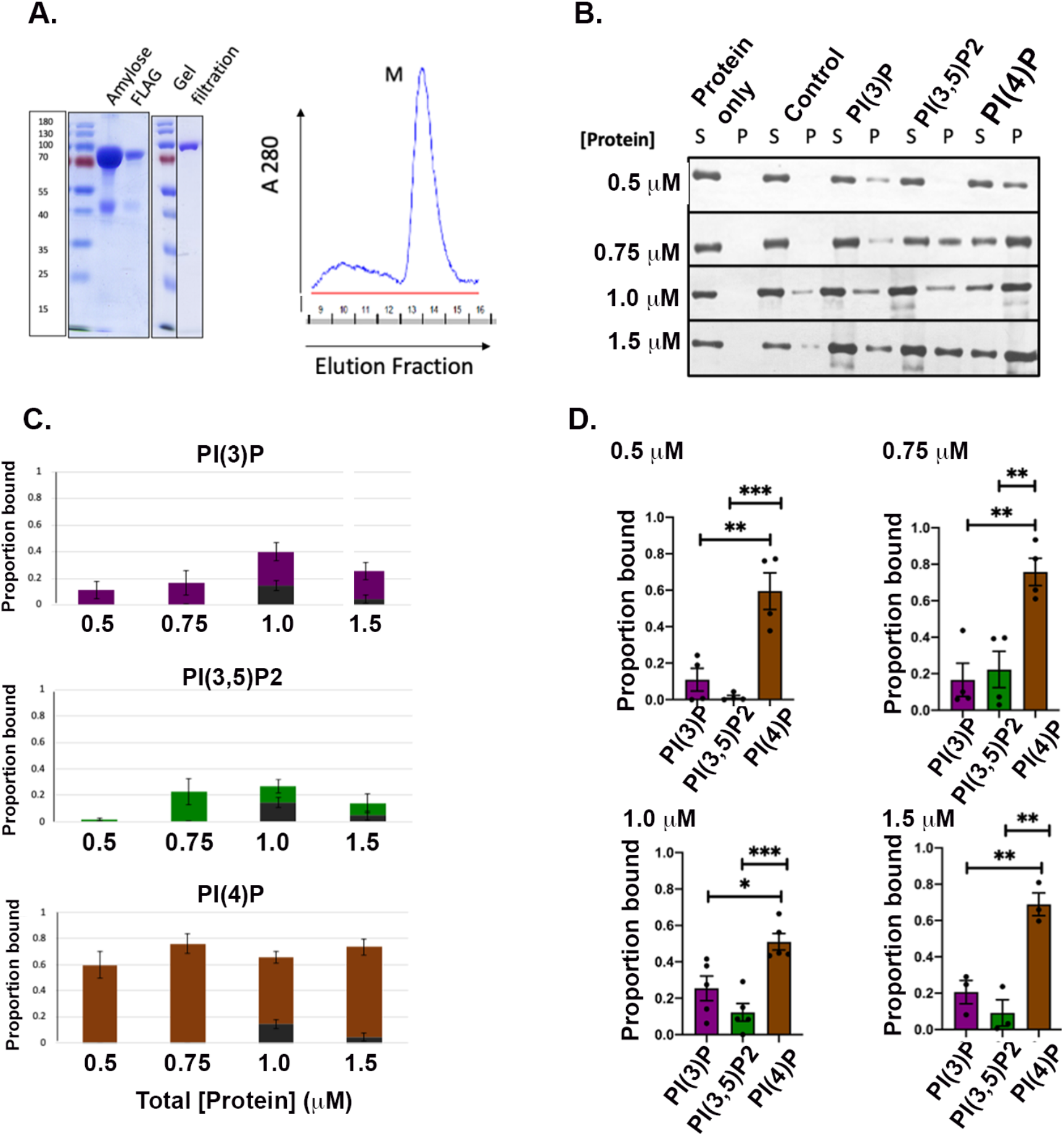
Hua2NT binds the Golgi PIP lipid PI(4)P *in vitro*. **A**. Left, Coomassie-stained SDS-PAGE showing purified MBP-Hua2NT-FLAG after amylose, FLAG column and gel filtration purification, with molecular mass markers. Right, Elution profile of MBP-Hua2NT-FLAG from gel filtration column. The peak corresponding to the monomeric molecular mass (M) was collected and used for liposome pelleting assay. **B**. Anti-FLAG immunoblot of MBP-Hua2NT-FLAG of supernatant and pellet fractions from protein only, control, and experimental liposome pelleting assays performed as described in Figure 2. **C**. Stacked plots of the mean proportion of protein bound to control liposomes (gray, bottom) vs. that specific to PIP-containing liposomes (colored, top). Error bars indicate S.E.M. **D**. Proportion of PIP-specific binding of MBP-Hua2NT-FLAG to liposomes containing the indicated lipids at different protein concentrations. At least three assays were performed for each protein concentration and PIP lipid, and each dot represents a distinct biological replicate. Band intensities were determined by ImageJ. Statistical significance was determined by ordinary one-way ANOVA with Tukey’s multiple comparison test. **** P<0.0001, *** P<0.0005, ** P<0.005, * P<0.05, ns P>0.05.

### Identification of candidate PIP lipid binding sites on human a1NT and a2NT

Once the binding specificities of Hua1NT and Hua2NT isoforms for different PIP lipids were established, we wanted to identify candidate PIP lipid binding sites in these isoforms. Identifying these binding sites will aid in understanding the functions of PIP lipids *in vivo* and recognizing PIP binding codes on V-ATPase subunit isoforms might aid in understanding regulation and targeting of V-ATPases. Many cytosolic or peripheral membrane proteins bind PIP lipids with an orthodox binding site like PX, PHD or FYVE (Strahl and Thorner 2007, Hammond and Balla 2015). None of these canonical binding sites are present in yeast or human V-ATPase a-subunit isoforms. However, some membrane can bind PIPs through noncanonical binding sites. To identify candidate PIP lipid binding sites, we used structure and homology modeling to identify aNT mutations for testing and then tested the mutant proteins for altered PIP lipid binding. The cryo-EM structure of human V-ATPase containing the a1 isoform from HEK293F cells (PDB 6WM2) has been reported (Wang, Wu et al. 2020). There is no structure for human V-ATPases containing the a2 isoform, so we used the molecular threading software PHYRE2 (Kelley, Mezulis et al. 2015) to model Hua2NT on the Hua1NT structure. An overlay of the a1NT structure and a2NT model is depicted in Figure 4A. The two subunit isoforms are moderately conserved (48.2% identity) (Supporting information 1). To identify candidate PIP lipid binding sites, we applied three major criteria. These include: (a) minimal homology between isoforms that show distinct PIP binding specificities, (b) presence of basic amino acid residues, capable of ionic interactions with negatively charged PIP head groups (possibly flanked by aromatic amino acids), and (c) predicted orientation of the binding sites toward the cytoplasmic leaflet of membrane, where the PIP lipids reside, in the assembled V-ATPase complex. In Hua1NT, Y^214^VH and G^239^FR were identified as candidate binding sequences (Figure 4B, left) that satisfied the criteria and in Hua2NT K^221^WY and G^183^KV were identified (Figure 4B, right). All of these residues are present in the distal domain in the loops in Hua1NT and Hua2NT. Although yeast Stv1NT has a binding site in a proximal loop, the corresponding proximal loops Hua1NT and Hua2NT are shorter and have no basic residues.

**Figure 4:**
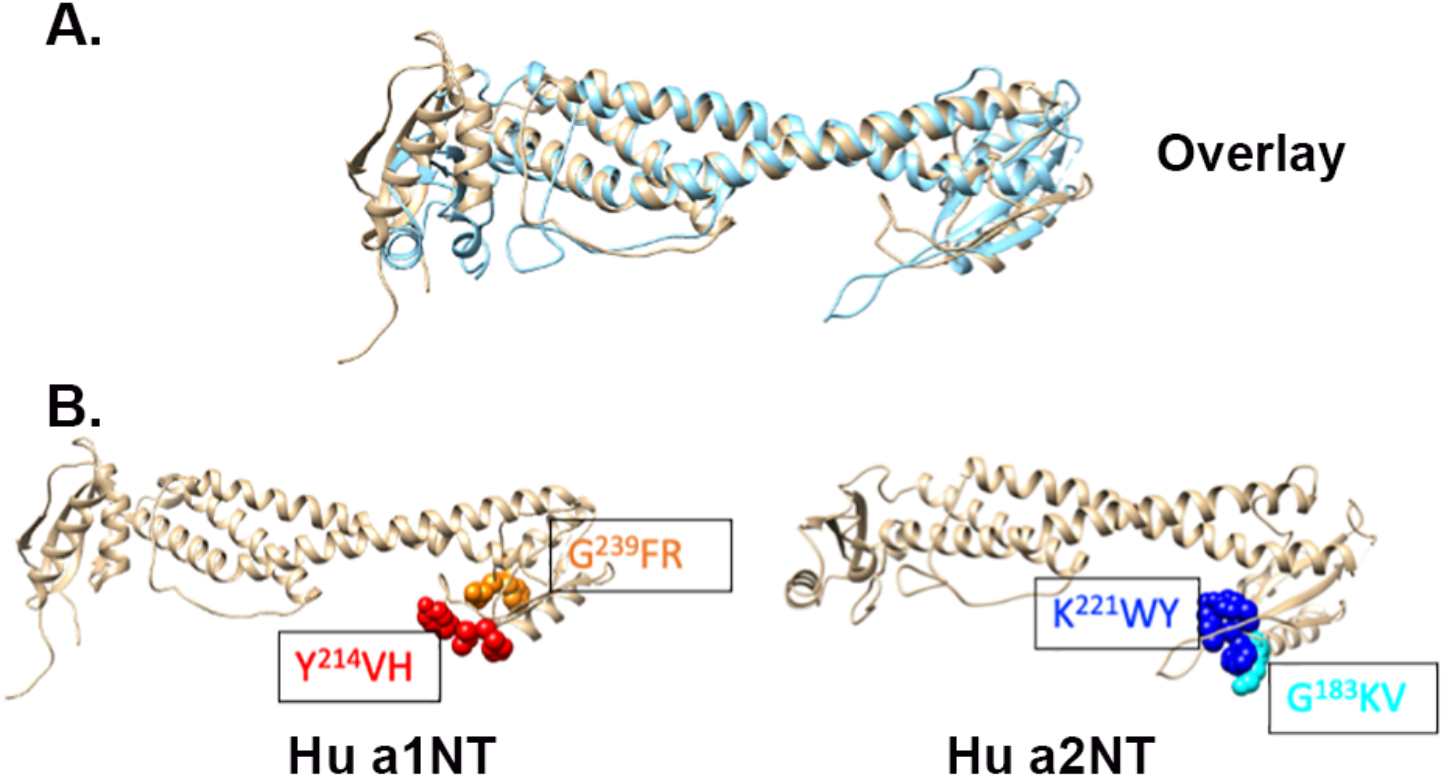
Identification of candidate PIP lipid binding sites on human a1NT and a2NT. **A**. Overlay of ribbon diagrams for the Hua1NT structure and Hua2NT model. The gold ribbon represents Hua1NT, and the light blue ribbon represent Hua2NT. **B**. Left, Hua1NT candidate PIP lipid binding sites Y^214^VH and G^239^FR with side chains shown in red and orange. Right, candidate PIP lipid binding sites of Hua2NT, K^221^WY and G^183^KV with side chains shown in blue and cyan.

### Hua1NT requires Y^214^VH sequence to bind PI(3)P and PI(3,5)P_2_

The candidate binding sites identified in Hua1NT, Y^214^VH and G^239^FR, were mutated using the primers listed in Table 1, to yield MBP-Hua1NT(Y^214^VH-AVA)-FLAG and MBP-Hua1NT(G^239^FR-AAA)-FLAG fusion proteins. The purified mutant proteins had similar purification profiles to the wild-type Hua1NT (Figure 5A, left), and the monomer fractions (M) from gel filtration (Figure 5A, right) were again used for liposome pelleting assay. Since wild-type Hua1NT showed binding specificity to liposomes containing PI(3)P and PI(3,5)P_**2**_, we tested the mutant proteins for binding to liposomes containing these PIP lipids. The immunoblots show that Hua1NT with Y^214^VH – AVA mutation has reduced binding to liposomes containing PI(3)P and PI(3,5)P_**2**_ as depicted by the more intense bands in supernatant (S) (Figure 5B, left) and the lower proportion of bound protein (Figure 5B, right). However, Hua1NT with G^239^FR – AAA mutation still binds to the liposomes containing PI(3)P and PI(3,5)P_**2**_ as indicated by bands in the pellets (P) (Figure 5C, right) and the relatively high proportion of protein bound to liposomes containing both lipids (Figure 5C-left). The PIP specific binding to both PI(3)P and PI(3,5)P_**2**_ of the wild-type and mutant versions of Hua1NT are compared in Figure 5D. The graph shows that Hua1NT with mutation at Y^214^VH site has almost no detectable binding for PI(3)P or PI(3,5)P_**2**_ and Hua1NT with mutation at G^239^FR site has slightly reduced binding to both lipids at low concentrations, suggesting a somewhat lower affinity, but binds similarly to wild-type at higher concentrations (Figure 5D). These data indicate that Hua1NT requires the Y^214^VH sequence to bind PI(3)P and PI(3,5)P2 *in vitro*.

**Table 1:**
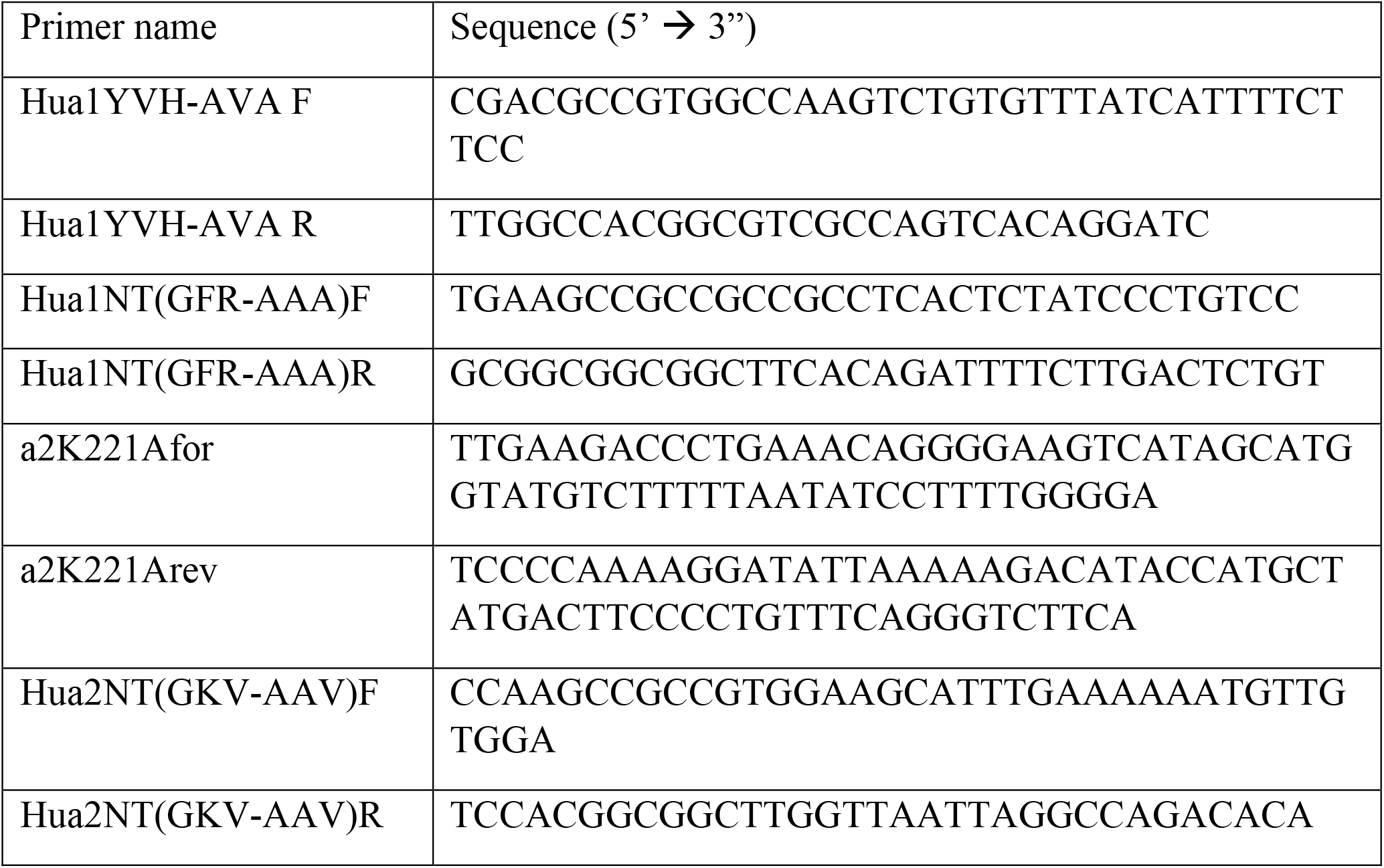
Oligonucleotides used in this study

**Figure 5:**
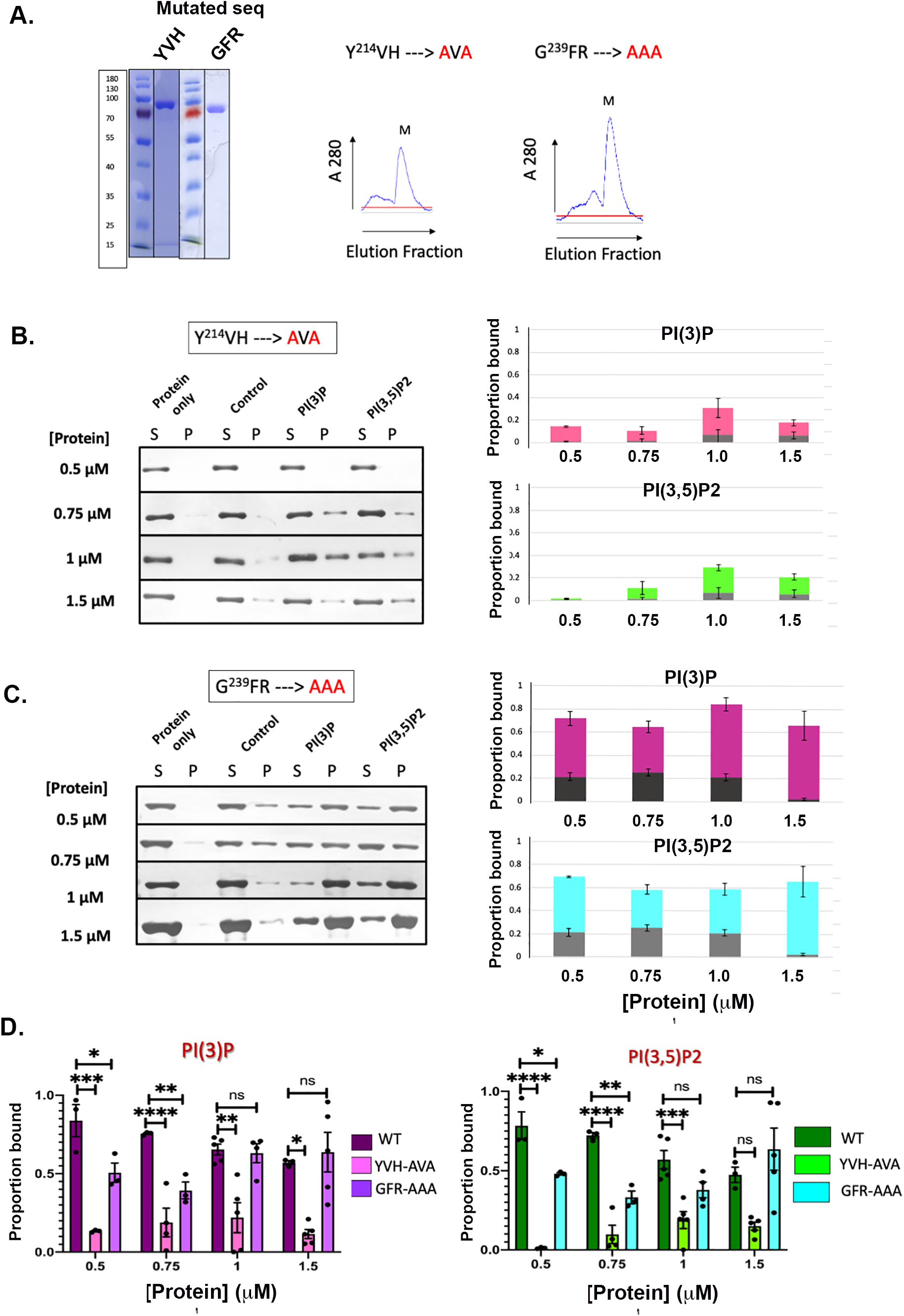
Hua1NT requires Y^214^VH sequence to bind PI(3)P and PI(3,5)P_2_. **A**. Left, Coomassie-stained SDS-PAGE showing purified MBP-Hua1(Y^214^VH -AVA)NT-FLAG and MBP-Hua1(G^239^FR -AAA)NT-FLAG after gel filtration. Right, Elution profiles of the two mutant proteins. The peaks corresponding to the monomeric molecular mass (M) were collected and used in the liposome pelleting assay. **B**. Left, Anti-FLAG immunoblot of MBP-Hua1(Y^214^VH -AVA)NT-FLAG at different concentrations with supernatant (S) and Pellet (P) fractions collected from protein only and control samples as described in Figure 2 or from liposomes 5% of PI(3)P or PI(3,5)P2 as indicated. Right, stacked plots showing the proportion of protein in the pellet in control samples (gray) or from samples containing the indicated PIP lipid (color) as described in Figure 2. **C**. Left, Anti-FLAG immunoblot of MBP-Hua1(G^239^FR - AAA)NT-FLAG at different concentrations in supernatant (S) and Pellet (P) fractions collected from pelleting with liposomes containing 5% of the PIP lipids. Right, stacked plot indicating the proportion of total protein bound to control (gray) vs. PIP specific binding (colored portion) at each protein concentration. Error bars represent S.E.M. **D**. Comparison of the proportion bound to PI(3)P (left) and PI(3,5)P2 (right) for each mutant to the wild-type protein shown in Figure 2. At least three assays were performed for each concentration and PIP lipid and each dot represents a distinct biological replicate. Comparison was done by ordinary one-way ANOVA with Tukey’s multiple comparison test. **** P<0.0001, *** P<0.0005, **

### Hua2NT requires K^221^WY sequence to bind PI(4)P

Interestingly, yeast Stv1, which resides in the Golgi, contains a high affinity binding site for PI(4)P in a W^83^KY sequence at its proximal end; mutation of only the lysine in this sequence abolished PI(4)P binding (Banerjee and Kane 2017). Given the similarity to the K^221^WY sequence in the distal end of Hua2NT, we mutated the lysine of the K^221^WY sequence to alanine to yield MBP-Hua2NT(K^221^A)-FLAG using the primers listed in Table1. In addition, the G^183^KV sequence, was mutated to generate an MBP-Hua2NT(G^183^KV-AAV)-FLAG fusion protein. The mutated proteins had similar purification profiles to wild-type Hua2NT (Figure 6A). The monomer fractions (M) were used in liposome pelleting assays. Since wild-type Hua2NT showed binding specificity for PI(4)P, we tested the binding of the mutants to PI(4)P-containing liposomes. The immunoblots show that Hua2NT with K^221^A mutation binds poorly to PI(4)P-containing liposomes, and the low proportion of MBP-Hua2NT(K^221^A)-FLAG in pellets (P) over several experiments (Figure 6B). Comparison of the PIP specific binding of Hua2NT mutant (K^221^A) with binding of the wild-type Hua2NT to PI(4)P-containing liposomes (Figure 6C) indicates significantly reduced binding of the mutant protein. For MBP-Hua2NT(G^183^KV-AAV)-FLAG, the immunoblot and graph depicting proportion of protein bound (Figure 6D) show that, curiously, Hua2NT with mutation at G^183^KV sequence binds to PI(4)P lipids at low protein concentrations (0.5 µM), but seems to have reduced binding to PI(4)P at higher concentration of protein. This behavior is very different than that of Hua2NT with the K^221^A mutation or the wild-type and mutant forms of Stv1NT and Hua1NT and will require further analysis to understand. However, it is clear that mutation of Hua2NT K^221^ to A greatly decreases binding to PI(4)P.

**Figure 6:**
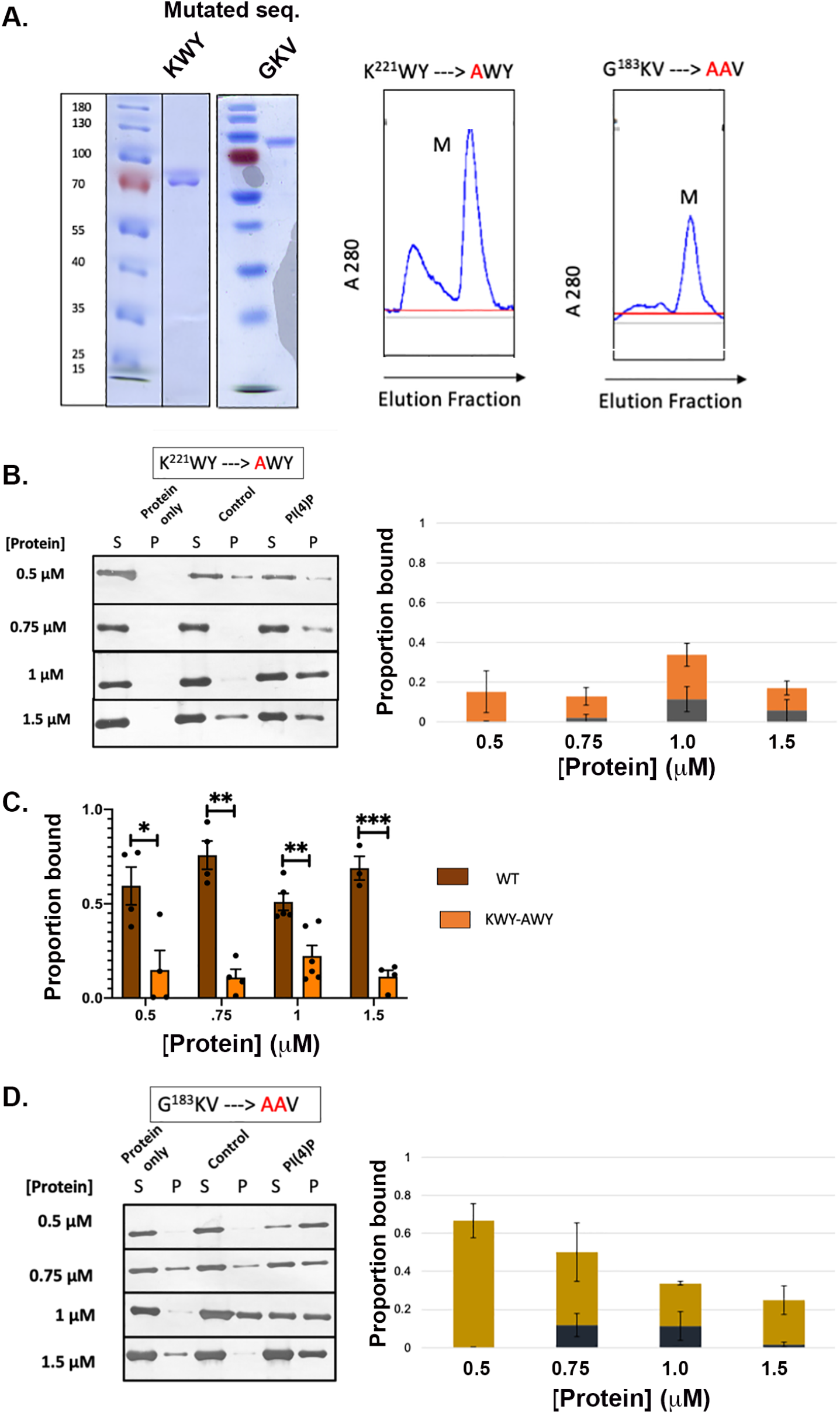
Hua2NT requires the sequence K^221^WY to bind PI(4)P. **A**. Left, Coomassie-stained SDS-PAGE showing purified MBP-Hua2(K^221^WY -AWY)NT-FLAG and MBP-Hua2(G^183^KV -AAV)NT-FLAG proteins with molecular mass marker. Right, Elution profiles of MBP-Hua2(K^221^WY -AWY)NT-FLAG and MBP-Hua2(G^183^KV -AAV)NT-FLAG from gel filtration column. The peaks corresponding to the monomeric molecular mass (M) were collected and used for liposome pelleting assays. **B**. Left, Anti-FLAG immunoblot of MBP-Hua2(K^221^WY -AWY)NT-FLAG in supernatant (S) and Pellet (P) fractions collected from protein only, control, and liposomes containing 5% PI(4)P. Total protein concentrations are indicated to the left of the blots. Right, Stacked plots showing the proportion of total protein in the control (gray) and the PI(4)P-specific proportion (orange). **C**. Comparison of PI(4)P-specific binding of wild-type Hua2NT (from Figure 3) to that of the mutant K^221^WY. Each dot represents a distinct biological replicate. Statistical significance was determined by ordinary one-way ANOVA with Tukey’s multiple comparison test. **** P<0.0001, *** P<0.0005, ** P<0.005, * P<0.05, ns P>0.05. **D**. Left, Anti-FLAG immunoblot of MBP-Hua2(G^183^KV -AAV)NT-FLAG at different concentrations in supernatant (S) and Pellet (P) fractions collected from controls or liposomes containing 5% PI(4)P. Right, Stacked plots of average proportion of total protein bound to control liposomes (gray) vs. PI(4)P-specific binding (gold).

### Different PIP binding specificities are encoded at a similar location in aNT isoform structures

These data highlight the importance of the Y^214^VH sequence in Hua1NT and K^221^WY in the Hua2NT sequence for binding to their respective PIP lipids. The sequence alignment in Figure 7A indicates that these two sequences are at very similar positions in the aligned sequences. A comparison of the three-dimensional structures indicates that both of the sequences lie in a poorly conserved loop at the distal end of the aNT isoforms. Remarkably, the sequence implicated in binding of yeast Vph1NT to PI(3,5)P_2_ is in a very similar position as shown in Figure 7B. This distal domain loop is poorly resolved in most V-ATPase cryo-EM structures, suggesting that it may be mobile, but is resolved one of the yeast (Khan, Lee et al. 2022) and some of the human V-ATPase structures (Wang, Wu et al. 2020, Wang, Long et al. 2020). Overall structure of V-ATPases is highly conserved between organisms and between complexes containing different a-subunit isoforms. These data suggest that there is a variable loop in the distal domain of multiple aNT isoforms that has the capacity for binding to PIP lipids. Intriguingly, different sequences at this location appear to support distinct PIP specificity.

**Figure 7:**
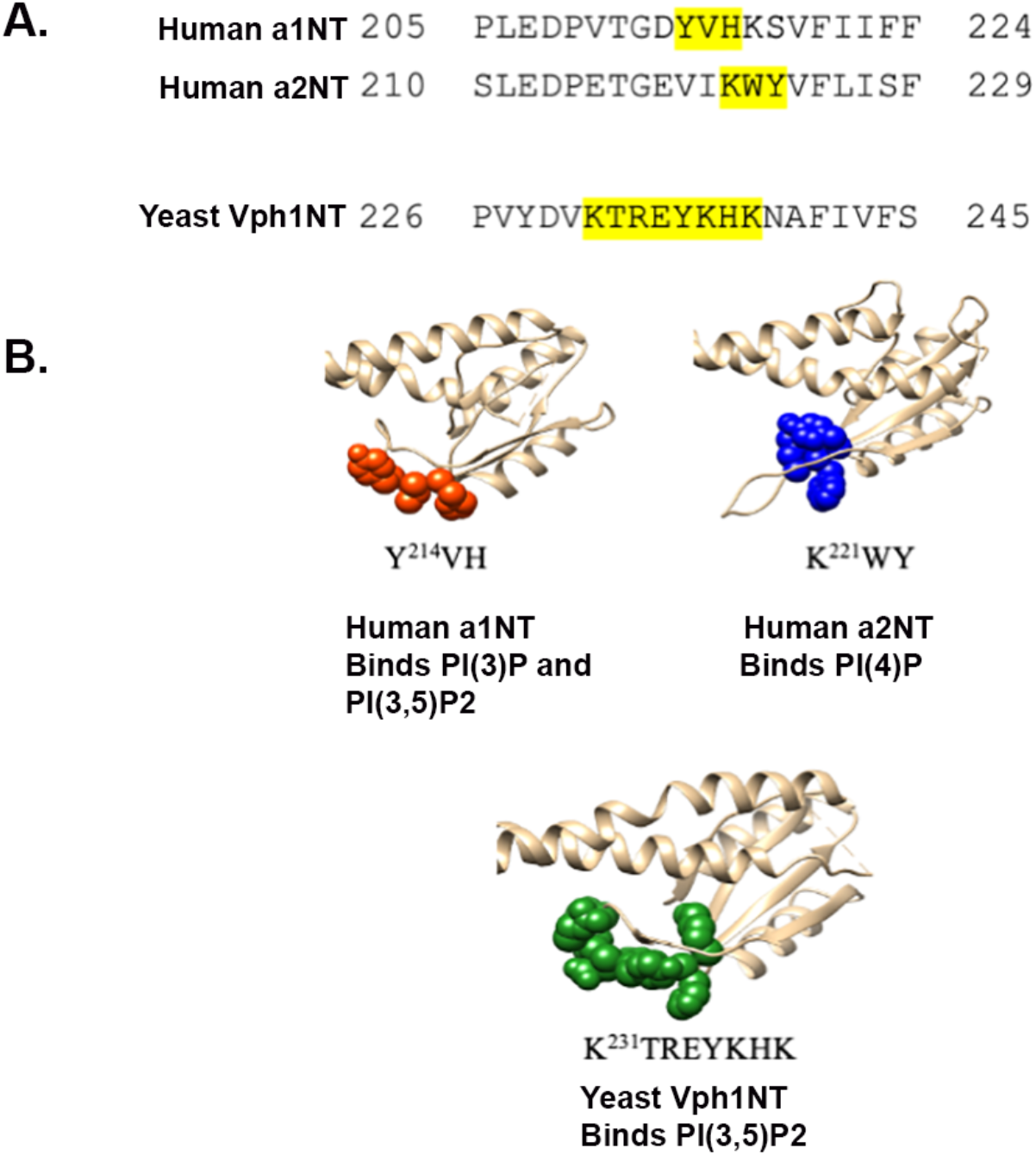
Different PIP binding specificities are encoded at a similar location in aNT isoform structures. **A**. Sequence alignment of a region in the distal end of Vph1NT, Hua1NT, and Hua2NT. Sequences that compromised binding to PIP lipids when mutated are highlighted. **B**. Ribbon diagram of distal end binding domains of Hua1NT, Hua2NT and Vph1NT showing the respective binding sequences, Y^214^VH (red), K^221^WY (blue), and K^231^TREYKHK (green).

## DISCUSSION

### PIP binding is a conserved feature of V-ATPase aNT domains

These data suggest that the capacity for PIP-specific binding is present in a-subunit isoforms from mammals as well as yeast, despite the lack of previously characterized PIP binding domains in any of the a-subunit isoforms. However, there are both similarities and differences in the binding of yeast and human isoforms to PIP lipids.

Human a2NT and yeast Stv1 are both primarily in Golgi-localized V-ATPases. Both bind tightly and specifically to liposomes containing PI(4)P, the PIP lipid enriched in the Golgi apparatus. The mutations that compromised PI(4)P binding, K^84^A in the W^83^KY of yeast Stv1NT and mutation of K^221^A in a K^221^WY sequence in human a2NT, appear to occur in a similar sequence context but are present at opposite ends of the aNT domain. Stv1 is longer than most a-subunit isoforms across species, and much of this additional length is in insertions in the NT domain, including a longer and more basic membrane-adjacent loop at the proximal end that contains the W^83^KY sequence. In the a2NT model, there are no basic amino acids in the shorter proximal loop. However, it is also notable that although mutation of Stv1 K^84^ abolishes binding to PI(4)P liposomes *in vitro* and transfers PI(4)P binding to Vph1NT when added at a similar position, experiments with chimeras between regions of Stv1NT and Vph1NT suggest that the distal end of Stv1NT may also help support high affinity binding to PI(4)P (Tuli and Kane 2023). The sequences involved are not known, and it is notable that Stv1 has a glutamate, which could disrupt PIP binding, in the region aligned with the K^221^WY sequence of a2NT. However, a recent study identified another distal end sequence conserved across yeast and human a-subunit isoforms that supports binding to several different PIP lipids, but has limited specificity *in vitro* (Chu, Yao et al. 2023). This sequence could help to strengthen PI(4)P binding in both Stv1NT and a2NT. In yeast, PI(4)P binding is linked to both activation of V-ATPase activity *in vitro* and retention or retrieval of Stv1-V-ATPases to the Golgi apparatus *in vivo*. Introduction of the K^221^A mutation into full-length human a2 will be necessary to determine whether the functional effects of PI4P binding are similar.

Comparisons of the human a1NT and yeast Vph1NT, both of which reside in endolysosomal compartments, reveals sequence conservation between the regions that compromise PI(3,5)P_2_ binding *in vitro* when mutated (Figure 7). In yeast, mutations in either of two adjacent sequences K^231^TREY- and K^236^HK-AAA block PI(3,5)P_2_ activation of V-ATPases in isolated vacuoles and cause defects in PI(3,5)P_2_-dependent osmotic stress responses (Banerjee, Clapp et al. 2019). Interestingly, the human Y^214^VH sequence implicated in binding of Hua1NT to PI(3,5)P_2_ aligns with a Y^235^KH sequence in the middle of the two sequences implicated in yeast Vph1NT binding to PI(3,5)P_2_ and both are followed by an additional lysine that was not mutated in this study, but could potentially contribute to binding (Figure 7). However, there are important differences as well. Although human a1NT binds tightly and specifically to liposomes containing either PI(3)P or PI(3,5)P_2_, Vph1NT binds poorly to liposomes *in vitro* (Tuli and Kane 2023), suggesting a lower affinity for PIP lipids than Hua1NT. In addition, V-ATPase activity in yeast could be activated by PI(3,5)P_2_ but not by PI(3)P, suggesting the two lipids have distinct effects on activity, and potential differences in binding (Banerjee, Clapp et al. 2019). Although similar activity assays are not yet available for the human a1 isoform, it is difficult to justify differences in specificity and affinity purely from the sequences in this region, as Vph1NT has additional basic amino acids including a lysine in the middle of the homologous sequence that could potentially contribute to binding to either or both of these PIP lipids (Figure 7). In addition, the basic lysine and arginine of the KTREY sequence are not present in a1NT. Functionally, the V-ATPase in the yeast vacuole appears to be closely integrated with the osmotic stress response, which is characterized by dramatic changes in PI(3,5)P_2_ levels in yeast (Duex, Nau et al. 2006, Li, Diakov et al. 2014). In this context, the rather weak affinity of Vph1NT for PI(3,5)P_2_ may be important for a robust regulatory response to the rapid rise of PI(3,5)P_2_ with osmotic stress. Although PI(3,5)P_2_ is also extremely important in the endolysosomal system of higher eukaryotes, the changes in lipid levels with environmental stress are less dramatic (Jin, Jin et al. 2017), so higher affinity binding could be appropriate to maintain both a basal level of interaction between V-ATPases and the PIP lipids support the response to stress. Notably both Vph1NT and Hua1NT are likely to transit through the Golgi en route to endosomes and lysosomes, but neither binds significantly to PI(4)P. This is potentially consistent with PIP binding occurring when they reach their compartments of residence, possibly accompanied by activation or other functional effeccts. Further experiments are necessary to establish how PI(3,5)P2 and PI(3)P affect function of V-ATPases containing the a1 isoform in cells.

### How could PIP binding sites in the distal domain of aNTs affect V-ATPase structure and activity?

As described above, it is remarkable that the human a1NT and a2NT show distinct PIP-specific binding that can be disrupted by mutations in sequences mapping to such as similar structural location (Figure 7A). We have found that binding to PI(3,5)P_2_ activates V-ATPases containing the Vph1 isoform, and also stabilizes them against disassembly of the V_1_ subcomplex from V_o_(Li, Diakov et al. 2014). The evidence that Vph1NT is regulated by sequences at this same location suggests that PIP binding to the mammalian a-subunit isoform to this region could also be functionally important. Structures are available for both intact yeast V-ATPases and isolated V_2_ subcomplexes containing Vph1, and we compared the position of the loop containing the potential PI(3,5)P_2_ binding sequence in both (Figure 8). Both the proximal and distal domains of Vph1NT collapse toward the center of the V_o_ subcomplex in the absence of V_1_(Stam and Wilkens 2017, Roh, Stam et al. 2018). In this conformation, the loop implicated in PI(3,5)P2 binding is above the c-ring and does not appear to be readily accessible to membrane lipids. In contrast, Vph1NT is in a more extended conformation in the assembled Vph1-containing V-ATPases where it now would have access to membrane lipids (Zhao, Benlekbir et al. 2015, Khan, Lee et al. 2022). In this extended conformation, Vph1NT interacts with V_1_ subunits in two of the three peripheral stalks, as required for stable V-ATPase assembly. The distal end of Vph1NT, specifically, interacts with one EG stalk and the foot end of the C subunit (Oot and Wilkens 2012, Oot, Couoh-Cardel et al. 2017). A similar extended conformation of Hua1NT is seen in structures of the intact V-ATPases containing this subunit (Abbas, Wu et al. 2020, Wang, Wu et al. 2020, Wang, Long et al. 2020). We hypothesize that PIP binding to the extended conformation of aNT helps to stabilize the assembled conformation. Reversible disassembly of V_1_ from V_o_, possibly initiated by departure of V_1_ subunit C, is a characteristic feature of V-ATPases from multiple species (Oot, Couoh-Cardel et al. 2017). It is possible that the V-ATPase is poised for disassembly even in its assembled state, and stabilizing factors, such as PIP binding are able to favor or stabilize the assembled conformation.

**Figure 8:**
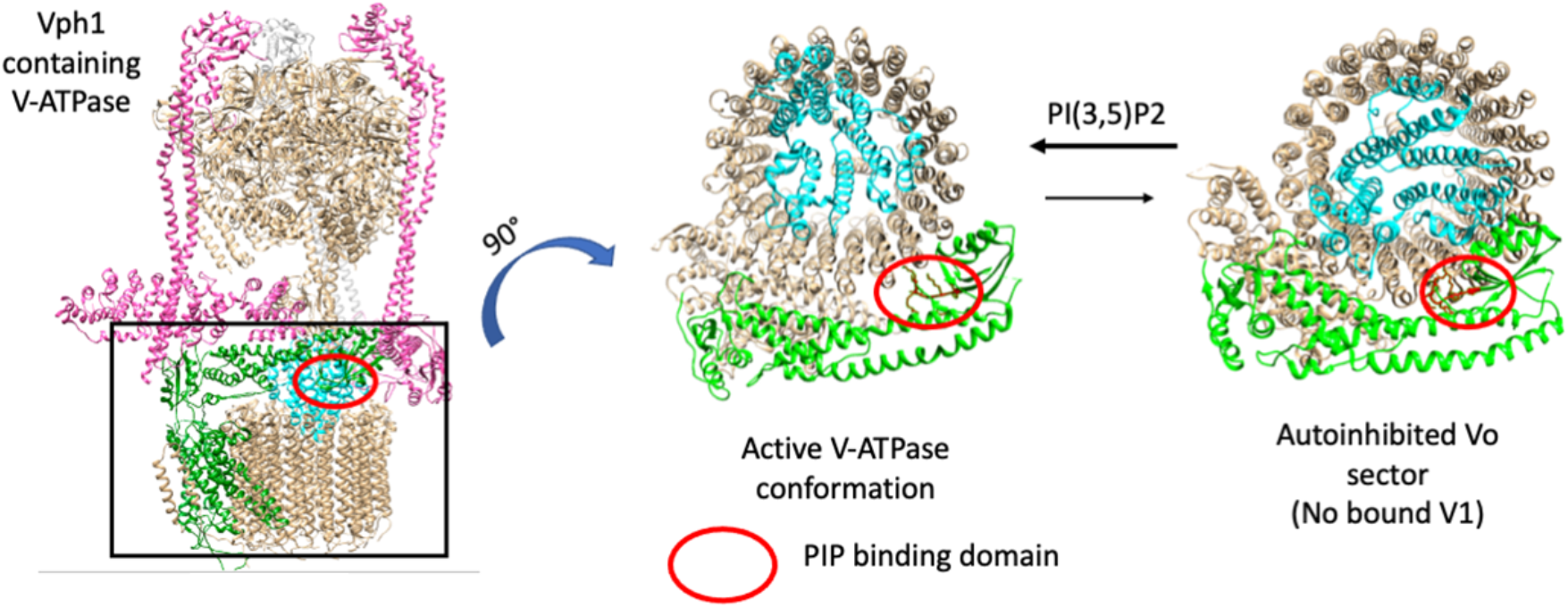
Position of the PI(3,5)P_2_ binding sequence in the assembled yeast V-ATPase and disassembled Vo sector: Comparison of position of PI(3,5)P2 binding sequence (circled red) in assembled V-ATPase and autoinhibited Vo sector, both containing the Vph1 isoform. The figures have been adopted from PDB 7FDA for the fully assembled complex (Khan et al., 2022) and PDB 6C6L or the isolated Vo complex (Roh et al., 2020).

There is still much to learn about the interaction between PIP lipids and V-ATPases, although this work indicates that the capacity for these interactions is conserved and focuses attention on the distal end of aNT isoforms as sites of PIP binding and regulation. The PIP binding mutations in Hua1NT and Hua2NT identified here can be introduced into the intact a-subunit isoforms to assess the functional importance of PIP binding. Mutation of the endogenous isoform genes may be necessary, given the presence of multiple a-subunit isoforms and the potential for overexpression to perturb isoform distribution or introduce functional compensation. In addition, although cryo-EM structures of V-ATPases have identified lipids within the V_2_ subcomplex, there is as yet no structure that shows direct interaction of the aNT distal end with lipid headgroups. Achieving such a structure may require restoration of PIP lipids after solubilization of V-ATPases, with the goal of preserving PIP interactions through reconstitution.

## EXPERIMENTAL PROCEDURES

### Cloning, expression, and purification of HuaNT isoforms from *E. coli*

MBP-Hua1NT(1-356)-FLAG fusion protein was generated by Anne Smardon (SUNY Upstate Medical University). PCR fragments containing the first 356 amino acids from human cDNA ATP6Voa1 were cloned into pGEM and excised with EcoRI and SalI, then cloned into pMAL:PPase cut with the same enzymes. Construction of MBP-Hua2NT-FLAG domain was described (Banerjee and Kane 2017). The mutant constructs, MBP-Hua1(Y^214^VH-AVA)NT-FLAG, MBP-Hua1(G^239^FR-AAA)NT-FLAG, MBP-Hua2(K^221^WY -AWY)NT-FLAG and MBP-Hua2(G^183^KV-AAV)NT-FLAG were constructed by introducing point mutations using In-Fusion HD Cloning (Takara Bio USA, Inc) with the primers listed in Table 1. The constructs were transformed into Rosetta [genotype: F− *ompT hsdS*B(rB− mB−) *gal dcm* (DE3) pRARE (CamR)] (Novogen) competent *E. coli* cells and grown to a density of A600 of 0.6 in 2.5% Luria-Bertani (LB) broth (Fisher Bioreagents) with 125 μg/ml ampicillin (Sigma-Aldrich) and 34 μg/ml chloramphenicol(Sigma-Aldrich). Expression of protein was then induced by addition of 0.3 mM isopropyl β-d-1-thiogalactopyranoside followed by incubation at 19°C for 16 hours. The cells were pelleted at 3000 rpm at 4°C for 20 minutes and resuspended in 25 ml amylose column buffer (20mM Tris-HCl, pH 7.4, 0.2M NaCl and 1mM EDTA). 2 mM MgCl_2_, 1mg/ml lysozyme, 2μg/ml DNase, 5 mM Dithiothreitol (DTT), 1 mM of Phenyl Methane Sulfonyl Fluoride (PMSF) were added, and the mixture was sonicated on ice for cell lysis, followed by slow rocking at 4°C for 30 minutes. The lysate was centrifuged at 15,000 rpm at 4°C for 20 mins. The supernatant was diluted 1:4 with the same buffer and purified through 5 ml Amylose and 1 ml FLAG affinity purification columns sequentially. Amylose resin was purchased from New England BioLabs and anti-FLAG M2 resin from Sigma. Purification in the amylose column involved running the lysate twice through the column followed by a wash with 10 column volumes of the same buffer. Proteins were eluted from amylose column with 10mM maltose monohydrate. 1 ml fractions were collected and 5 mM DTT was added. Peak fractions with protein concentrations giving an A_2_> 0.5 were pooled and dialyzed in dialysis tubing with MWCO 3.5kD (Spectrum Labs) in FLAG buffer (50mM Tris-HCl, pH 7.4 and 150mM NaCl) overnight at 4°C. The dialyzed mixture was added to an anti-FLAG M2 column (Sigma), for the second round of purification. The column was washed with 10 column volumes of FLAG buffer and proteins were eluted as 1 ml fractions with 100 μg/ml FLAG peptide (Sigma). 5 mM DTT was added to each fraction and peak fractions as determined from Coomassie blue gel electrophoresis were pooled and concentrated in Vivaspin 15-max speed filter (MWCO 30kD) (Sartorius Stedim Lab Ltd.) to a final concentration of at least 1mg/ml. The concentrated protein was run on a Superdex Increase 200 (Cytivia) gel-filtration FPLC column on Biologic DuoFlow System (Biorad). The fraction corresponding to the peak corresponding to the molecular mass of the monomer protein (∼80kDa) was collected and was used for liposome pelleting assay.

### Liposome preparation and liposome pelleting assay

Lipids were purchased from Avanti Polar Lipids as lyophilized powder and dissolved in CHCl_3_: CH_3_ OH: H_2_ O in 20:9:1 ratio. Liposomes with 50 mol % 16:0 phosphatidylcholine (PC), 25 mol % 16:0 phosphatidylserine (PS), 18 mol %16:0 phosphatidylthanolamine (PE), 2 mol % 16:0 NBD-PE (for visualizing liposomes) and 5 mol % 18:1 phosphatidylinositol phospholipids PI(3)P, PI(4)P or PI(3,5)P2, were prepared by in Liposome making buffer (50mM Tris, 25mM NaCl, pH 7.4) by extrusion as described by Banerjee and Kane, 2017 (Banerjee and Kane 2017). Control liposomes, without PIP lipids were prepared with an extra 5 mol% 16:0 PS.

Liposome pelleting assays were performed by incubating the prepared liposomes with the expressed protein of interest at room temperature for 30 minutes to allow protein-lipid interaction. The salt concentration was adjusted to 60 mM with NaCl. The final volume of the mixture was 100 µl and samples contained protein concentrations of 0.5, 0.75, 1 and 1.5 µM with a final lipid concentration of 0.33 mM. Following incubation, the mixture was centrifuged at 95,000 rpm at 4°C for 30 mins in a TLA-100 fixed angle rotor using Beckman Coulter mini ultracentrifuge. The supernatant was collected, and the pellet was resuspended in 100 µl buffer with 50mM Tris-HCl and 60mM NaCl, pH 7.4. Both supernatant and pellet samples were precipitated with 10% trichloro acetic acid (TCA), and the resulting pellets were washed with cold acetone and resuspended in 30 µl cracking buffer (50mM Tris-HCl, pH 6.8, 8M urea, 5% SDS and 1mM EDTA). Equal volumes of each sample were separated by SDS-PAGE and transferred into nitrocellulose membrane. The blots were probed with mouse monoclonal anti-FLAG M2 antibody (Sigma) and anti-mouse IgG alkaline phosphatase conjugated antibody (Promega). Band intensities were quantified by Image J. To calculate proportion of total protein that bound to the liposomes with PIP or without PIP (control), the band intensities of the pellet fractions were divided by the total band intensity of the supernatant and pellet. The PIP specific binding was calculated by subtracting the band intensity of PIP independent binding (control) from band intensity of PIP specific binding. Freshly prepared liposomes and purified proteins were used in the experiments.

### Protein structure prediction

The homology model of Hua2NT (1-364) was generated by PHYRE2 (Protein Fold Recognition server -One-to-One threading software) (Kelley, Mezulis et al. 2015) from the structure of V-ATPase from human embryonic kidney cell line HEK293F (PDB 6WM2) (Wang, Wu et al. 2020). A 100% confidence model of the submitted sequence with 88% coverage based on rat brain rotational state 2 bound to ADP and Sidk (PDB c6vq7a) (Abbas, Wu et al. 2020). NCBI-BLASTp program was used to align the protein sequences of the two HuaNT isoforms and identify regions that are not conserved between isoforms and have positively charged basic amino acids with aromatic (and hydrophobic) amino acids. Structures were visualized on UCSF Chimera Version 1.16 (Pettersen, Goddard et al. 2004).

### Statistical Analysis

All experiments were performed at least three times for each protein concentration and PIP lipid. ImageJ was used to quantify the bands in the immunoblots. Data is represented as mean ±SEM. Significance was determined using Ordinary one-way ANOVA with multiple comparison in GraphPad Prism9. p values: *, p < 0.05, **, p < 0.005, ***, p < 0.0005 and ****, p < 0.00005.

## ACKNOWLEDGEMENTS

This work was supported by NIH R01 GM126020 and R35 GM14256 to P.M.K. The authors thank Anne Smardon and Subhrajit Banerjee for plasmid constructions, Stephan Wilkens, Rebecca Oot, and the Kane lab for helpful discussions, and Wenyi Feng for suggestions on data analysis and presentation.

## REFERENCES

Abbas, Y. M., D. Wu, S. A. Bueler, C. V. Robinson and J. L. Rubinstein (2020). “Structure of V-ATPase from the mammalian brain.” Science 367(6483): 1240–1246.

Arata, Y., T. Nishi, S. Kawasaki-Nishi, E. Shao, S. Wilkens and M. Forgac (2002). “Structure, subunit function and regulation of the coated vesicle and yeast vacuolar (H+)-ATPases.” Biochimica et Biophysica Acta (BBA)-Bioenergetics 1555(1-3): 71–74.

Balla, T. (2013). “Phosphoinositides: tiny lipids with giant impact on cell regulation.” Physiological reviews 93(3): 1019–1137.

Banerjee, S., K. Clapp, M. Tarsio and P. M. Kane (2019). “Interaction of the late endo-lysosomal lipid PI (3, 5) P2 with the Vph1 isoform of yeast V-ATPase increases its activity and cellular stress tolerance.” Journal of Biological Chemistry 294(23): 9161–9171.

Banerjee, S., K. Clapp, M. Tarsio and P. M. Kane (2019). “Interaction of the late endo-lysosomal lipid PI(3,5)P2 with the Vph1 isoform of yeast V-ATPase increases its activity and cellular stress tolerance.” J Biol Chem 294(23): 9161–9171.

Banerjee, S. and P. M. Kane (2017). “Direct interaction of the Golgi V-ATPase a-subunit isoform with PI (4) P drives localization of Golgi V-ATPases in yeast.” Molecular biology of the cell 28(19): 2518–2530.

Banerjee, S. and P. M. Kane (2020). “Regulation of V-ATPase Activity and Organelle pH by Phosphatidylinositol Phosphate Lipids.” Front Cell Dev Biol 8: 510.

Breton, S. and D. Brown (2013). “Regulation of luminal acidification by the V-ATPase.” Physiology (Bethesda) 28(5): 318–329.

Chandra, M., Y. K.-Y. Chin, C. Mas, J. R. Feathers, B. Paul, S. Datta, K.-E. Chen, X. Jia, Z. Yang and S. J. Norwood (2019). “Classification of the human phox homology (PX) domains based on their phosphoinositide binding specificities.” Nature communications 10(1): 1528.

Chu, A., Y. Yao, G. T. Saffi, J. H. Chung, R. J. Botelho, M. Glibowicka, C. M. Deber and M. F. Manolson (2023). “Characterization of a PIP Binding Site in the N-Terminal Domain of V-ATPase a4 and Its Role in Plasma Membrane Association.” International Journal of Molecular Sciences 24(5): 4867.

Chu, A., R. A. Zirngibl and M. F. Manolson (2021). “The V-ATPase a3 Subunit: Structure, Function and Therapeutic Potential of an Essential Biomolecule in Osteoclastic Bone Resorption.” Int J Mol Sci 22(13).

Clague, M. J., S. Urbé and J. de Lartigue (2009). “Phosphoinositides and the endocytic pathway.” Experimental cell research 315(9): 1627–1631.

Collins, M. P. and M. Forgac (2020). “Regulation and function of V-ATPases in physiology and disease.” Biochim Biophys Acta Biomembr 1862(12): 183341.

De Craene, J.-O., D. L. Bertazzi, S. Bär and S. Friant (2017). “Phosphoinositides, major actors in membrane trafficking and lipid signaling pathways.” International journal of molecular sciences 18(3): 634.

Dove, S. K., K. Dong, T. Kobayashi, F. K. Williams and R. H. Michell (2009). “Phosphatidylinositol 3, 5-bisphosphate and Fab1p/PIKfyve underPPIn endo-lysosome function.” Biochemical Journal 419(1): 1–13.

Duex, J. E., J. J. Nau, E. J. Kauffman and L. S. Weisman (2006). “Phosphoinositide 5-phosphatase Fig 4p is required for both acute rise and subsequent fall in stress-induced phosphatidylinositol 3,5-bisphosphate levels.” Eukaryot Cell 5(4): 723–731.

Forgac, M. (2007). “Vacuolar ATPases: rotary proton pumps in physiology and pathophysiology.” Nature reviews Molecular cell biology 8(11): 917–929.

Frattini, A., P. J. Orchard, C. Sobacchi, S. Giliani, M. Abinun, J. P. Mattsson, D. J. Keeling, A. K. Andersson, P. Wallbrandt, L. Zecca, L. D. Notarangelo, P. Vezzoni and A. Villa (2000). “Defects in TCIRG1 subunit of the vacuolar proton pump are responsible for a subset of human autosomal recessive osteopetrosis.” Nat Genet 25(3): 343–346.

Graham, L. A., B. Powell and T. Stevens (2000). “Composition and assembly of the yeast vacuolar H (+)-ATPase complex.” Journal of Experimental Biology 203(1): 61–70.

Hammond, G. R. and T. Balla (2015). “Polyphosphoinositide binding domains: Key to inositol lipid biology.” Biochimica Et Biophysica Acta (BBA)-Molecular and Cell Biology of Lipids 1851(6): 746–758.

Hammond, G. R., M. J. Fischer, K. E. Anderson, J. Holdich, A. Koteci, T. Balla and R. F. Irvine (2012). “PI4P and PI(4,5)P2 are essential but independent lipid determinants of membrane identity.” Science 337(6095): 727–730.

Hille, B., E. J. Dickson, M. Kruse, O. Vivas and B.-C. Suh (2015). “Phosphoinositides regulate ion channels.” Biochimica Et Biophysica Acta (BBA)-Molecular and Cell Biology of Lipids 1851(6): 844–856.

Ho, C. Y., T. A. Alghamdi and R. J. Botelho (2012). “Phosphatidylinositol-3, 5-bisphosphate: no longer the poor PIP2.” Traffic 13(1): 1–8.

Hurtado-Lorenzo, A., M. Skinner, J. El Annan, M. Futai, G. H. Sun-Wada, S. Bourgoin, J. Casanova, A. Wildeman, S. Bechoua, D. A. Ausiello, D. Brown and V. Marshansky (2006). “V-ATPase interacts with ARNO and Arf6 in early endosomes and regulates the protein degradative pathway.” Nat Cell Biol 8(2): 124–136.

Jaskolka, M. C., S. R. Winkley and P. M. Kane (2021). “RAVE and Rabconnectin-3 Complexes as Signal Dependent Regulators of Organelle Acidification.” Front Cell Dev Biol 9: 698190.

Jin, N., Y. Jin and L. S. Weisman (2017). “Early protection to stress mediated by CDK-dependent PI3,5P2 signaling from the vacuole/lysosome.” J Cell Biol 216(7): 2075–2090.

Kane, P. M. (2006). “The where, when, and how of organelle acidification by the yeast vacuolar H+-ATPase.” Microbiology and Molecular Biology Reviews 70(1): 177–191.

Kane, P. M. (2006). “The where, when, and how of organelle acidification by the yeast vacuolar H+-ATPase.” Microbiol Mol Biol Rev 70(1): 177–191.

Kawasaki-Nishi, S., K. Bowers, T. Nishi, M. Forgac and T. H. Stevens (2001). “The amino-terminal domain of the vacuolar proton-translocating ATPase a subunit controls targeting and in vivo dissociation, and the carboxyl-terminal domain affects coupling of proton transport and ATP hydrolysis.” Journal of Biological Chemistry 276(50): 47411–47420.

Kelley, L. A., S. Mezulis, C. M. Yates, M. N. Wass and M. J. Sternberg (2015). “The Phyre2 web portal for protein modeling, prediction and analysis.” Nature protocols 10(6): 845–858.

Khan, M. M., S. Lee, S. Couoh-Cardel, R. A. Oot, H. Kim, S. Wilkens and S. H. Roh (2022). “Oxidative stress protein Oxr1 promotes V-ATPase holoenzyme disassembly in catalytic activity-independent manner.” EMBO J 41(3): e109360.

Kohio, H. P. and A. L. Adamson (2013). “Glycolytic control of vacuolar-type ATPase activity: a mechanism to regulate influenza viral infection.” Virology 444(1-2): 301–309.

Kornak, U., E. Reynders, A. Dimopoulou, J. van Reeuwijk, B. Fischer, A. Rajab, B. Budde, P. Nurnberg, F. Foulquier, A. D.-t. S. Group, D. Lefeber, Z. Urban, S. Gruenewald, W. Annaert, H. G. Brunner, H. van Bokhoven, R. Wevers, E. Morava, G. Matthijs, L. Van Maldergem and S. Mundlos (2008). “Impaired glycosylation and cutis laxa caused by mutations in the vesicular H+-ATPase subunit ATP6V0A2.” Nat Genet 40(1): 32–34.

Li, S. C., T. T. Diakov, T. Xu, M. Tarsio, W. Zhu, S. Couoh-Cardel, L. S. Weisman and P. M. Kane (2014). “The signaling lipid PI (3, 5) P2 stabilizes V1–Vo sector interactions and activates the V-ATPase.” Molecular biology of the cell 25(8): 1251–1262.

Li, S. C., T. T. Diakov, T. Xu, M. Tarsio, W. Zhu, S. Couoh-Cardel, L. S. Weisman and P. M. Kane (2014). “The signaling lipid PI(3,5)P(2) stabilizes V(1)-V(o) sector interactions and activates the V-ATPase.” Molecular Biology of the Cell 25(8): 1251–1262.

Maxson, M. E. and S. Grinstein (2014). “The vacuolar-type H(+)-ATPase at a glance -more than a proton pump.” Journal of Cell Science 127(Pt 23): 4987–4993.

Maxson, M. E. and S. Grinstein (2014). “The vacuolar-type H+-ATPase at a glance–more than a proton pump.” Journal of cell science 127(23): 4987–4993.

McCartney, A. J., Y. Zhang and L. S. Weisman (2014). “Phosphatidylinositol 3,5-bisphosphate: low abundance, high significance.” Bioessays 36(1): 52–64.

Nishi, T. and M. Forgac (2000). “Molecular cloning and expression of three isoforms of the 100-kDa a subunit of the mouse vacuolar proton-translocating ATPase.” J Biol Chem 275(10): 6824–6830.

Ochotny, N., A. Van Vliet, N. Chan, Y. Yao, M. Morel, N. Kartner, H. P. von Schroeder, J. N. Heersche and M. F. Manolson (2006). “Effects of human a3 and a4 mutations that result in osteopetrosis and distal renal tubular acidosis on yeast V-ATPase expression and activity.” Journal of Biological Chemistry 281(36): 26102–26111.

Oot, R. A., S. Couoh-Cardel, S. Sharma, N. J. Stam and S. Wilkens (2017). “Breaking up and making up: The secret life of the vacuolar H+ -ATPase.” Protein Sci 26(5): 896–909.

Oot, R. A., S. Couoh-Cardel, S. Sharma, N. J. Stam and S. Wilkens (2017). “Breaking up and making up: The secret life of the vacuolar H+-ATPase.” Protein Science 26(5): 896–909.

Oot, R. A. and S. Wilkens (2012). “Subunit interactions at the V1-Vo interface in yeast vacuolar ATPase.” J Biol Chem 287(16): 13396–13406.

Pamarthy, S., A. Kulshrestha, G. K. Katara and K. D. Beaman (2018). “The curious case of vacuolar ATPase: regulation of signaling pathways.” Mol Cancer 17(1): 41.

Pettersen, E. F., T. D. Goddard, C. C. Huang, G. S. Couch, D. M. Greenblatt, E. C. Meng and T. E. Ferrin (2004). “UCSF Chimera—a visualization system for exploratory research and analysis.” Journal of computational chemistry 25(13): 1605–1612.

Qi, J. and M. Forgac (2008). “Function and subunit interactions of the N-terminal domain of subunit a (Vph1p) of the yeast V-ATPase.” Journal of Biological Chemistry 283(28): 19274–19282.

Roh, S.-H., N. J. Stam, C. F. Hryc, S. Couoh-Cardel, G. Pintilie, W. Chiu and S. Wilkens (2018). “The 3.5-Å CryoEM structure of nanodisc-reconstituted yeast vacuolar ATPase Vo proton channel.” Molecular cell 69(6): 993–1004. e1003.

Roh, S. H., N. J. Stam, C. F. Hryc, S. Couoh-Cardel, G. Pintilie, W. Chiu and S. Wilkens (2018). “The 3.5-A CryoEM Structure of Nanodisc-Reconstituted Yeast Vacuolar ATPase Vo Proton Channel.” Mol Cell 69(6): 993–1004 e1003.

Seidel, T. (2022). “The Plant V-ATPase.” Front Plant Sci 13: 931777.

Shewan, A., D. J. Eastburn and K. Mostov (2011). “Phosphoinositides in cell architecture.” Cold Spring Harbor perspectives in biology 3(8): a004796.

Stam, N. J. and S. Wilkens (2017). “Structure of the Lipid Nanodisc-reconstituted Vacuolar ATPase Proton Channel: DEFINITION OF THE INTERACTION OF ROTOR AND STATOR AND IMPLICATIONS FOR ENZYME REGULATION BY REVERSIBLE DISSOCIATION.” J Biol Chem 292(5): 1749–1761.

Stam, N. J. and S. Wilkens (2017). “Structure of the lipid nanodisc-reconstituted vacuolar ATPase proton channel: definition of the interaction of rotor and stator and implications for enzyme regulation by reversible dissociation.” Journal of Biological Chemistry 292(5): 1749–1761.

Strahl, T. and J. Thorner (2007). “Synthesis and function of membrane phosphoinositides in budding yeast, Saccharomyces cerevisiae.” Biochimica et Biophysica Acta (BBA)-Molecular and Cell Biology of Lipids 1771(3): 353–404.

Stransky, L., K. Cotter and M. Forgac (2016). “The Function of V-ATPases in Cancer.” Physiol Rev 96(3): 1071–1091.

Sun-Wada, G.-H., H. Tabata, N. Kawamura, M. Aoyama and Y. Wada (2009). “Direct recruitment of H+-ATPase from lysosomes for phagosomal acidification.” Journal of cell science 122(14): 2504–2513.

Sun-Wada, G. H. and Y. Wada (2022). “The a subunit isoforms of vacuolar-type proton ATPase exhibit differential distribution in mouse perigastrulation embryos.” Sci Rep 12(1): 13590.

Toei, M., R. Saum and M. Forgac (2010). “Regulation and isoform function of the V-ATPases.” Biochemistry 49(23): 4715–4723.

Toyomura, T., T. Oka, C. Yamaguchi, Y. Wada and M. Futai (2000). “Three subunit a isoforms of mouse vacuolar H+-ATPase: preferential expression of the a3 isoform during osteoclast differentiation.” Journal of Biological Chemistry 275(12): 8760–8765.

Tuli, F. and P. M. Kane (2023). “Chimeric a-subunit isoforms generate functional yeast V-ATPases with altered regulatory properties in vitro and in vivo.” Mol Biol Cell: mbcE22070265.

Vasanthakumar, T., S. A. Bueler, D. Wu, V. Beilsten-Edmands, C. V. Robinson and J. L. Rubinstein (2019). “Structural comparison of the vacuolar and Golgi V-ATPases from Saccharomyces cerevisiae.” Proceedings of the National Academy of Sciences 116(15): 7272–7277.

Vasanthakumar, T. and J. L. Rubinstein (2020). “Structure and Roles of V-type ATPases.” Trends in biochemical sciences 45(4): 295–307.

Wallroth, A. and V. Haucke (2018). “Phosphoinositide conversion in endocytosis and the endolysosomal system.” Journal of Biological Chemistry 293(5): 1526–1535.

Wang, L., D. Wu, C. V. Robinson, H. Wu and T.-M. Fu (2020). “Structures of a complete human V-ATPase reveal mechanisms of its assembly.” Molecular cell 80(3): 501–511. e503.

Wang, L., D. Wu, C. V. Robinson, H. Wu and T. M. Fu (2020). “Structures of a Complete Human V-ATPase Reveal Mechanisms of Its Assembly.” Mol Cell 80(3): 501–511 e503.

Wang, R., T. Long, A. Hassan, J. Wang, Y. Sun, X. S. Xie and X. Li (2020). “Cryo-EM structures of intact V-ATPase from bovine brain.” Nat Commun 11(1): 3921.

Zhao, J., S. Benlekbir and J. L. Rubinstein (2015). “Electron cryomicroscopy observation of rotational states in a eukaryotic V-ATPase.” Nature 521(7551): 241–245.

