## Supplemental Figure 1 for "Human V-ATPase a-subunit isoforms bind specifically to distinct phosphoinositide phospholipids"

**Mitra and Kane, Supplemental Figure 1: Alignment of Human a1NT and a2NT**

```
Query  1    MGELFRSEEMTLAQLFLQSEAAYCCVSELGELGKVQFRDLNPDVNVFQRKFVNEVRRCEE  60
            MG LFRSE M LAQLFLQS  AY C+S LGE G VQFRDLN +V+ FQRKFV EV+RCEE
Sbjct  1    MGSLFRSETMCLAQLFLQSGTAYECLSALGEKGLVQFRDLNQNVSSFQRKFVGEVKRCEE  60

Query  61   MDRKLRFVEKEIRKANIPIMDTGENPEVPFPRDMIDLEANFEKIENELKEINTNQEALKR  120
            ++R L ++ +EI +A+IP+ +   +P  P  + +++++   +K+E EL+E+  N+E L++
Sbjct  61   LERILVYLVQEINRADIPLPEGEASPPAPPLKQVLEMQEQLQKLEVELREVTKNKEKLRK  120

Query  121  NFLELTELKFILRKTQQFFDE-AELHHQQMADPDLLEESSSLLEPSEMGRGTPLRLGFVA  179
            N LEL E   +LR T+ F     E        P L  ES SLL+ S M R    +LGFV+
Sbjct  121  NLLELIEYTHMLRVTKTFVKRNVEFEPTYEEFPSL--ESDSLLDYSCMQR-LGAKLGFVS  177

Query  180  GVINRERIPTFERMLWRVCRGNVFLRQAEIENPLEDPVTGDYVHKSVFIIFFQGDQLKNR  239
            G+IN+ ++  FE+MLWRVC+G   +  AE++  LEDP TG+ +   VF+I F G+Q+ ++
Sbjct  178  GLINQGKVEAFEKMLWRVCKGYTIVSYAELDESLEDPETGEVIKWYVFLISFWGEQIGHK  237

Query  240  VKKICEGFRASLYPCPETPQERKEMASGVNTRIDDLQMVLNQTEDHRQRVLQAAAKNIRV  299
            VKKIC+ +   +YP P T +ER+E+  G+NTRI DL  VL++TED+ ++VL  AA+++
Sbjct  238  VKKICDCYHCHVYPYPNTAEERREIQEGLNTRIQDLYTVLHKTEDYLRQVLCKAAESVYS  297


Query  300  WFIKVRKMKAIYHTLNLCNIDVTQKCLIAEVWCPVTDLDSIQFALRRGTEHSGSTVPSIL  359
              I+V+KMKAIYH LN+C+ DVT KCLIAEVWCP  DL  ++ AL  G+  SG+T+PS +
Sbjct  298  RVIQVKKMKAIYHMLNMCSFDVTNKCLIAEVWCPEADLQDLRRALEEGSRESGATIPSFM  357

Query  360  NRMQTNQTPPTYNKTNKFTYGFQNIVDAYGIGTYREIN
            N + T +TPPT  +TNKFT GFQNIVDAYG+G+YRE+N
Sbjct  358  NIIPTKETPPTRIRTNKFTEGFQNIVDAYGVGSYREVN
```
